# Surveillance of Japanese Encephalitis Virus in Piggery Effluent and Environmental Samples: A Complementary Tool for Outbreak Detection

**DOI:** 10.1101/2025.03.27.645166

**Authors:** Warish Ahmed, Metasebia Gebrewold, David T. Williams, Jianning Wang, Wendy J.M. Smith, Leah G. Starick, Regina Fogarty, Kirsty Richards, Stuart L. Simpson

## Abstract

Japanese encephalitis virus (JEV) is an emerging public health and biosecurity concern in Australia, with recent human cases and detections in mosquitoes and pigs across multiple states highlight the risk to susceptible human and animal populations. While traditional surveillance methods such as mosquito trapping, sentinel chicken programs and direct testing of pig specimens remain essential, monitoring effluent offers a valuable complementary approach for detecting infections within animal populations. This study presents the first evidence of JEV in Australian piggery effluents/environmental waters, demonstrating the feasibility of effluent and environmental water surveillance for JEV monitoring. Effluent/environmental samples from multiple piggery sites were analyzed using real-time reverse transcription polymerase chain reaction (RT-PCR), revealing the presence of JEV genetic fragments in solid and liquid fractions of effluents at three farms, with corresponding veterinary cases in some herds. Viral RNA was detected more frequently in solid fraction of effluent samples, aligning with previous findings on the partitioning behaviour of mosquito-borne viruses. The detection of JEV in the borrow pit (i.e., a man-made excavation that holds water) water sample highlights potential transmission pathways via mosquito vectors. These findings demonstrate the value of effluent monitoring as an additional tool for JEV surveillance in piggery settings, supporting potential early warning systems and mitigation strategies. Integrating effluent-based monitoring with traditional surveillance approaches could improve livestock industry related disease detection, risk assessments, and response efforts for human and animal health in endemic and emerging regions. Wastewater/effluent surveillance may have important applications for the management of a wide range of emerging animal diseases.

**IMPORTANCE:** This study presents the first evidence of Japanese encephalitis virus (JEV) detection in Australian piggery effluents, establishing effluent surveillance as a valuable complementary tool for monitoring viral pathogens in animal populations. Our findings support the integration of effluent monitoring with traditional surveillance systems to improve early warning capabilities, enhance biosecurity, and mitigate risks to both animal and human health.

## Introduction

Japanese encephalitis virus (JEV), a member of the genus *Orthoflavivirus* in the family *Flaviviridae*, represents a major etiological agent of epidemic encephalitis throughout the Asia and western Pacific nations (1-3). Following the emergence of JEV in south-eastern Australia in 2022, there is significant concern about continued re-emergence in seasons where favorable climatic conditions exist (2,4). JEV is maintained in an enzootic transmission cycle primarily involving ardeid wading birds and *Culex* mosquitoes (5-7). Wild birds and pigs serve as important amplifying hosts for JEV and pigs experience direct health impacts (i.e., reproductive disease) from the virus (7,8). The risk of epizootic spillover to humans has been associated with the farming of domestic pigs in endemic regions of southeast Asia (8).

To mitigate the growing impact of JEV transmission and its spread to new regions, disease management strategies include human and pig vaccination programs, mosquito control and bite mitigation measures, and public awareness and education (4,9). However, these combined measures may still be insufficient to fully control the spread, particularly in regions favouring mosquito proliferation with gaps in vaccination coverage. Many JEV endemic countries also do not routinely vaccinate domestic pigs. In Australia, there is currently no swine JEV vaccine approved for use. Continued research and enhanced surveillance efforts are therefore essential to effectively mitigate the risk of JEV. In this context, urban (human) wastewater surveillance can provide a population-level assessment of infection burden and has proven to be effective in detecting both symptomatic and asymptomatic cases of disease in communities across a range of settings (10; https://data.wastewaterscan.org/).

A prospective influenza A (IAV) RNA monitoring at 190 wastewater treatment plants (WWTPs) across the USA identified increases in IAV RNA concentrations at 59 plants coinciding with H5N1 outbreaks in dairy cattle. The H5 marker was found at four plants which were all located in affected states and receiving industrial discharges containing animal waste. These findings show that wastewater surveillance can detect animal-associated influenza signals and underscore the importance of accounting for agricultural inputs (11). As viral shedding often precedes veterinary symptoms, it enables earlier outbreak detection than traditional methods (12,13). Wastewater/effluent surveillance can be equally effective for detecting viral diseases in animal industry settings, where livestock population-level samples may be collected from waste streams in animal housing areas, wastewater/effluent management facilities (e.g., effluent pits and ponds), and environmental waters (14,15).

A case study of JEV detection in municipal wastewater during an outbreak highlights the potential of wastewater surveillance as a complementary tool to existing monitoring efforts (16). Expanding the scope of wastewater surveillance to include livestock effluents aligns with the principles of “One Health” and “Planetary Health,” providing valuable insights into the health of animal populations and connections between human, animal and environmental settings (17). Monitoring of viral pathogens like JEV in livestock population waste streams may contribute to early detection systems, which not only help manage human and animal population health but may also support economic stability where certain disease outbreaks threaten domestic and international trade resilience (15).

Experimental and field evidence has shown that domestic pigs shed JEV following infection (18-21). Detection of JEV nucleic acid and infectious virus has been reported in oronasal swabs for up to six and five days, respectively (18), and JEV RNA was detected in swine oral fluids for up to 11-24 days after inoculation (19,21). JEV genome has also been detected in the urine and feces of experimentally infected pigs (18,22). Virus could be detected in fecal swabs from 3 to 7 days post inoculation, with viral loads ranging from 32 to 213 genome equivalents per mL (22). These findings suggest that effluent surveillance in piggery environments could be effective, potentially offering early detection of JEV circulation and potentially identifying outbreaks before they escalate. This approach follows the successful approach of human wastewater surveillance in detecting viral pathogens.

No studies have yet detected JEV in piggery effluents, but the 2022 outbreak of JEV, together with the re-emergence in south-eastern Australia in 2025 (23) highlight the increasing risk of infection to human and animal health. Here, we present the first evidence of JEV in Australian piggery effluents. Our findings show that effluent and environmental monitoring can be a complementary tool for JEV surveillance in piggery settings, offering detection and monitoring at the animal population level. This proactive surveillance approach could complement traditional methods like mosquito monitoring, chicken sentinel surveillance and direct pig surveillance, enhancing overall detection, mitigation, and response strategies. The approach may have applications for the management of a wide range of emerging animal diseases.

## MATAERIALS AND METHODS

### Sampling and concentration of viruses

Effluent and environmental samples were collected from six pig farms (Farms 1–5 in Victoria and Farm 6 in Queensland, Australia) between 17/12/2024 and 27/02/2025 using a convenience sampling approach based on farm owner participation. Piggeries typically have a range of sheds designated for different pig categories or activities, e.g., sow, breeding, farrowing, gestating, and nursery etc. These provide opportunities for effluent monitoring before the effluent stream combines and mixes within pits and larger effluent ponds (Fig. 1). Effluent and environmental water samples were collected as below:

**FIG 1.**
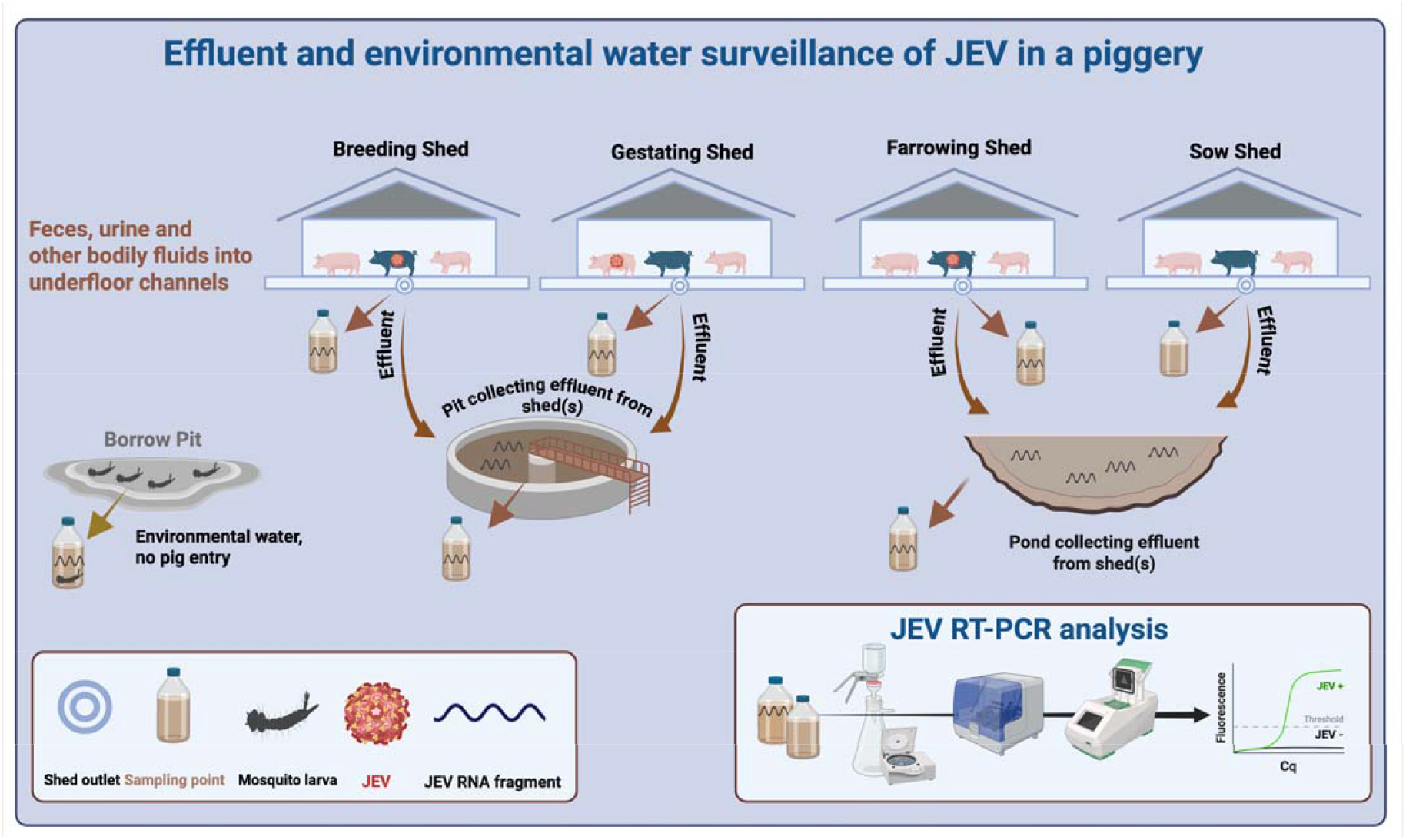
A typical piggery showing various sheds for pig housing, effluent systems and sampling points.

**Farm 1:** Four effluent grab samples (50 to 200 mL) were collected on different days from a pit receiving effluent from flush channels. Pits are in-ground tanks storing at least one day’s of effluent from multiple sheds.

**Farms 2-4:** Five, two, and one effluent sample were collected from effluent ponds located within 100 m of pig housing sheds for Farms 2, 3, and 4, respectively.

**Farm 5**: Four grab samples (50 to 200 mL) were collected from a pit, an effluent pond, farrowing shed, and breeding shed, plus one solid effluent sample from a sow shed.

**Farm 6:** Eight effluent grab samples (50 to 200 mL) were collected, with four from a gestating shed and four from an effluent pond on 12, 21, 24, and 27/02/2025. Additionally, three environmental water grab samples were collected from a borrow pit on 21, 24, and 27/02/2025. A borrow pit is a man-made excavation that holds water (no pig effluent) and was assumed to be a potential habitat for mosquito larvae. All samples were stored at 4°C up to 48 h before concentration and extraction.

### Viral DNA/RNA concentration from wastewater and nucleic acid extraction

An adsorption-extraction (AE) concentration method, with minor modifications for effluent liquid and solid fractions, was used to concentrate viruses from effluent samples (24). The AE workflow started with centrifuging 50 mL of an effluent sample at 4,750 *g* for 20 min. The supernatant was filtered through a 0.80-μm pore-size, electronegative hydroxyapatite (HA) membrane (47-mm diameter, AAWG04700) using a magnetic filter funnel (Pall Corporation, Port Washington, New York, USA) and a filter flask (Merck Millipore Ltd.), following the method described by Akter et al. (24). After filtration, the membrane was aseptically removed from the filter funnel, rolled, and placed into a 7-mL bead-beating tube with 1 g of 0.5 mm diameter Zirconium Oxide beads (Next Advance, Inc, NY, USA) for nucleic acid extraction. The pellet resulting from the centrifugation step was collected from the 50-mL tube and transferred into a 7-mL bead-beating tube for nucleic acid extraction. Water samples (∼50 to 100 mL) collected from the borrow pit were directly processed using the AE method without any centrifugation (24).

Nucleic acid extraction from the membrane (effluent liquid fraction) and the pellet (effluent solid fraction) was performed separately using the RNeasy PowerWater Kit (Cat. No. 14700– 50-NF, Qiagen). For each bead-beating tube containing the membrane or pellet, 990 μL of lysis buffer PM1 and 10 μL of β-mercaptoethanol (Cat. No. M6250–10 mL, Sigma-Aldrich) were added. The tube contents were then homogenized using a Precellys 24 tissue homogenizer (Bertin Technologies) at 9,000 rpm for three 15 s cycles, with a 10 s interval between cycles. After homogenization, the tubes were centrifuged at 4000 *g* for 5 min to pellet the filter debris, solids, and beads. Nucleic acid was eluted in 150 μL of DNase- and RNase-free water and stored at −20°C for 72 to 96 h prior to RT-PCR analysis.

### RT-PCR assays

The detection of JEV was carried out using RT-PCR assays targeting the NS-1 (Universal JEV assay) (25) and M (JEV G4 assay) genes of JEV. JEV G4 assay was designed by Australian centre for disease preparedness (ACDP), CSIRO. The Universal JEV assay enables the detection of all JEV genotypes, while the JEV-G4 assay specifically detects JEV genotype 4, the genotype currently circulating in Australia (2). RT-PCR assays were performed in two separate laboratories (Lab 1 and Lab 2). When a nucleic acid sample tested positive in Lab 1 (CSIRO Environment RU BC2 laboratory), it was subsequently sent to Lab 2 (Australian Centre for Disease Preparedness) for verification. This is due to JEV being classified as Category 1 restricted matter under the Queensland Biosecurity Act and consequently subject to certain testing and reporting requirements. This step was taken in accordance with the conditions outlined in the testing permit issued by Queensland Department of Agriculture and Fisheries (DAF), which stated that positive detections may require confirmation by a second laboratory. Initially, Universal JEV RT-PCR was conducted in Lab 1, but it was later replaced with the JEV G4 RT-PCR assay due to the latter’s increased sensitivity.

In Lab 1, positive controls were prepared from nucleic acid extracts of gamma irradiated JEV-G4. In Lab 1, Universal JEV RT-PCR amplifications were performed with a 20 µL reaction mixture containing 5 µL of TaqMan™ Fast Virus 1-Step Master Mix (Applied Biosystems, Thermo Fisher Scientific, MA, USA), 400 nM of forward primer (GCC ACC CAG GAG GTC CTT), 400 nM of reverse primer (CTT CCT CTC AGA ACC CCT ATC C) and 400 nM of probe (FAM-CTA ATG GAA CGC ATC CC-MGB). A 5 μL nucleic acid sample was used in each reaction. JEV G4 RT-PCR amplifications were performed with a 20 µL reaction mixture containing 5 µL of TaqMan™ Fast Virus 1-Step Master Mix (Applied Biosystems), 900 nM of forward primer (AAG AAG CCT GGC TGG ATT CA), 900 nM of reverse primer (GGA TTC CTT ATG ATC CAA TTT TCA G) and 250 nM of probe (FAM-CGA AAG CCA CCC GGT ATC TCA -MGB). A 5 μL nucleic acid sample was used in each reaction. Each run included triplicate positive controls and triplicate no-template controls. Universal JEV and JEV G4 assays amplified 63 and 73 bp of products, respectively. The RT-PCR assays were performed on a Bio-Rad CFX96 thermal cycler (Bio-Rad Laboratories, Hercules, California, USA) with manual settings for threshold and baseline (40 to 100 RFU). The RT-PCR cycling parameters were 10 min at 50°C, 10 min at 95°C, 45 cycles of 30 s at 95°C, and 45 s at 56°C for Universal JEV RT-PCR and 10 min at 50°C, 10 min at 95°C, 45 cycles of 30 s at 95°C, and 45 s at 60°C for G4 JEV RT-PCR. All RT-PCR reactions were performed in triplicate.

In Lab 2, JEV G4 RT-PCR amplifications were performed with a 15 µL reaction mixture containing 7.5 µL of AgPath-ID™ One-Step RT-PCR Reagents (Applied Biosystems), 900 nM of forward primer (AAG AAG CCT GGC TGG ATT CA), 900 nM of reverse primer (GGA TTC CTT ATG ATC CAA TTT TCA G) and 250 nM of probe (FAM-CTA ATG GAA CGC ATC CC-MGB). A 5 μL nucleic acid sample was used in each reaction. Each run included one positive control, one negative extraction control and one no-template control, each tested in duplicate. The RT-PCR assays were performed on a QuantStudio 5 real-time PCR systems (Thermo Fisher Scientific) with manual settings for threshold (0.1) and baseline (3-15). The RT-PCR cycling parameters were 10 min at 45°C, 10 min at 95°C, 45 cycles of 15 s at 95°C, and 45 s at 60°C for the G4 assay. All RT-PCR reactions were performed in duplicate.

### Quality assurance/controls

A SARS-CoV-2 RT-PCR assay was applied to determine inhibition in extracted nucleic acid samples from effluents by seeding known copy number (10^4^) of gamma-irradiated SARS-CoV-2 (CDC 2019-novel coronavirus (2019-nCoV) real-time RT-PCR diagnostic panel). The reference quantification cycle (Cq) value was determined from triplicate RT-PCR reactions containing only the positive control and compared with the Cq values obtained from all effluent nucleic acid samples. Samples were considered to have no inhibition when the Cq values of effluent nucleic acid samples were within 2 Cq values of the reference Cq value (26). Nucleic acid extraction and RT-PCR setup were performed in separate laboratories to minimize potential contamination.

### Sequencing

A 73 bp JEV G4 RT-PCR amplified product in one effluent sample from Farm 5 collected on 18/02/2025 was sequenced using a Sanger (Applied Biosystems, Foster City, CA, USA) sequencing platform to confirm JEV amplicon identity. This sample was chosen due to its low Cq value (30.2), indicating a high viral load suitable for sequencing. For Sanger sequencing, 5 µL of RT-PCR reaction was electrophoresed in a E-Gel™ SizeSelect™ II Agarose Gel, 2% and the E-Gel Power Snap Plus Electrophoresis System (Invitrogen™) following the manufacturers protocol for collecting size fragments. Target bands of expected size (i.e., 73 bp) were excised from the gel and submitted to a Sanger sequencing service provider along with forward and reverse primers (AGRF, Brisbane, Australia).

### Data interpretation

In Lab 1, samples were classified as positive for JEV by RT-PCR if amplification was detected in at least one out of three replicates within 40 cycles, whereas in Lab 2, samples were considered positive if amplification was detected in at least one of two replicates within the same cycle threshold. Samples were considered indeterminate when the Cq value was between 40 to 45. A sample was considered not-detected (ND) or negative when none of the three or two replicates showed amplification.

## RESULTS AND DISCUSSION

All RT-PCR negative controls consistently returned negative results, confirming the absence of contamination, while positive controls reliably tested positive for each run, ensuring RT-PCR assay validity. For RT-PCR inhibition assay, the reference Cq value was 30.8, and the Cq values for effluent nucleic acid samples ranged from 30.1 to 31.2. These values were well within the acceptable ±2 Cq value, indicating no inhibition in the nucleic acid samples. The piggery effluent samples were analyzed for the presence of JEV nucleic acid across various sampling locations within three farms (1, 5, and 6) over multiple sampling dates (Table 1).

**TABLE 1.**
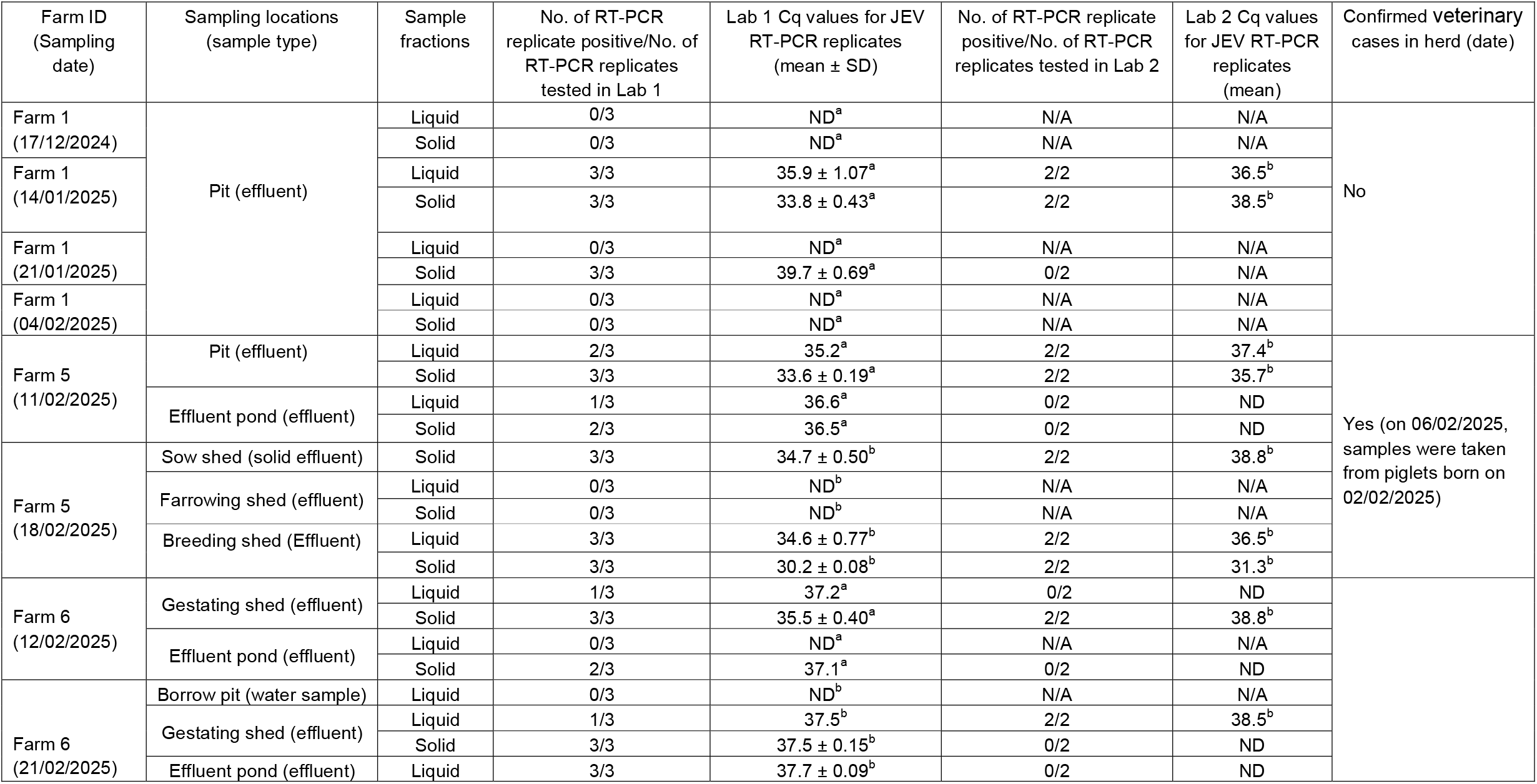
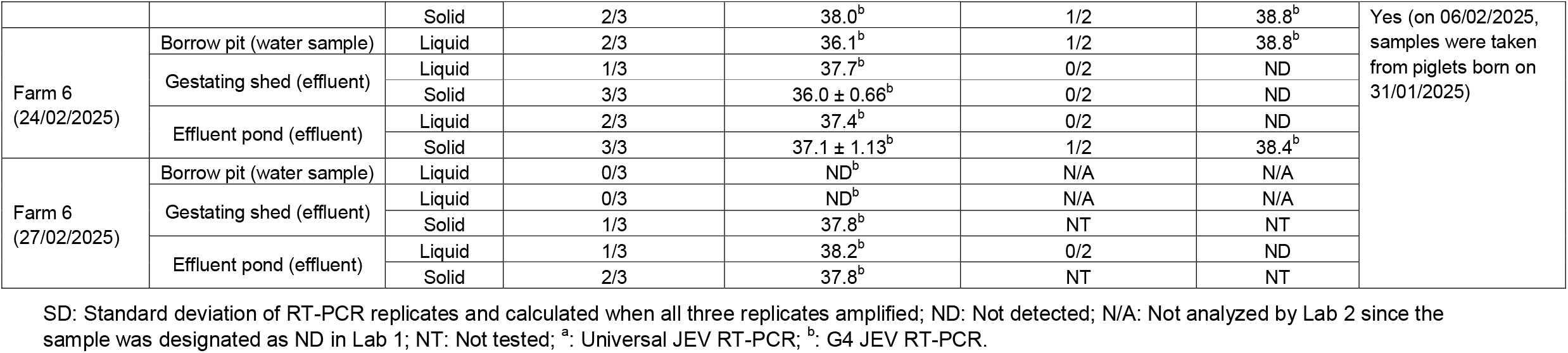
Detection of JEV in piggery effluent samples and environmental samples.

JEV nucleic acid was not detected in any effluent samples from Farm 2, Farm 3, and Farm 4, and no veterinary cases were reported in the herds on these farms. The absence of JEV nucleic acid in effluent samples suggests that the JEV was either not present or present at levels below the assay limit of detection (ALOD). The ALOD values for Universal JEV and JEV G4 assays were determined to be ∼6 copies/reaction (data not shown). The former scenario aligns with the lack of veterinary cases, suggesting that JEV transmission did not occur on these farms during the sampling period. At Farm 1, the Universal JEV assay did not detect JEV in either the liquid or solid fractions of effluent samples collected from the pit on 17/12/2024.The first detection of JEV occurred on 14/01/2025, in both liquid and solid fraction of effluent samples collected from the same pit. All three Universal JEV RT-PCR replicates tested positive at Lab 1. The mean Cq values were 35.9 ± 1.07 for liquid fraction samples and 33.8 ± 0.43 for solid fraction samples run on the same PCR plate, indicating a higher viral load in the solid fractions. Potentially corresponding to a 4-fold greater concentration, assuming ideal RT-PCR efficiency. This could suggest that viral particles are more effectively retained in solid fraction compared to liquid fraction. Our findings are in accordance with Roldan-Hernandez et al. (27) who reported that seeded mosquito borne viruses may partition several orders of magnitudes higher in solid fractions than in liquids in wastewater samples collected from the primary sludge line of wastewater treatment plants in the USA.

Follow-up sampling at Farm 1 on 21/01/2025 Universal JEV assay detected JEV only in the solid fraction of effluent (3/3 replicates positive, Cq = 39.7 ± 0.69), while the liquid fraction tested negative. By 04/02/2025 both matrices were RT-PCR negative for JEV. These combined results suggest a possible shift to detectable JEV levels only in solid fractions, followed by a potential decline in shedding or an effect of JEV dilution or potential decay of JEV genome in the sampling environments. It is also possible that the infected pigs ceased shedding or were shedding at levels that were below the ALOD. In this regard, RT-PCR detection and virus isolation studies indicated oronasal shedding of JEV can persist for between 5 to 6 days and can be detected from pen-based chew rope samples 11 to 24 days after virus inoculation (18,19,21). Fecal shedding of JEV in experimentally infected swine appeared from day three and persisted for three to seven days, with the average viral load ranging from 32 to 213 genome equivalents per mL (22). These findings suggest that the transient nature of JEV shedding in pigs may explain the decline in detection over time observed in this study. Interestingly, Farm 1 reported no veterinary cases of JE, despite the detection of viral nucleic acid in effluent samples, indicating that asymptomatic pigs were likely contributing to viral shedding. Farm 1 is a breeding farm comprising sows and newborn piglets (up to 3 to 4 weeks old). Natural infection of pigs is usually clinically inapparent, and typically manifests as reproductive disease in pregnant sows, leading to abortion, stillbirths and mummified fetuses, which becomes apparent at farrowing (8,28). In young pigs, JEV infection can occasionally cause a wasting syndrome with neurological signs. No veterinary JE cases have been confirmed on Farm 1 as of 27/02/2025. However, as not all sows pregnant during the collection of JEV positive effluent samples had farrowed, and given the potential for vertical transmission or past-farrowing infections, future cases cannot be ruled out.

Effluent samples collected from Farm 5 on 11/02/2025 tested positive for JEV using the Universal JEV assay. Liquid fraction sample from the pit showed 2/3 positive replicates (mean Cq = 35.2), while solid fraction sample was 3/3 positive replicates (mean Cq ± SD = 33.6 ± 0.19). A second effluent sample collected on the same day from an effluent pond also yielded positive results, albeit at lower detection levels, with liquid fraction samples showing 1/3 positive replicates (Cq = 36.6) and solid fraction samples 2/3 positive replicates (Cq = 36.5). Farm 5 had a confirmed JE veterinary case, with a litter of stillborn and mummified piglets born 02/02/2025 testing RT-PCR-positive for JEV in cerebrospinal fluid and placenta samples, aligning with the effluent RT-PCR detection results.

On 18/02/2025, three additional samples were collected from Farm 5 from separate breeding, farrowing and sow sheds. A solid effluent sample from the sow shed tested positive in all three replicates (Cq: 34.7 ± 0.50) using the JEV G4 assay. Lab 2 also confirmed the result, detecting JEV in 2/2 replicates (Cq: 38.8) using the JEV G4 assay. Effluent samples from the farrowing shed showed no JEV detection in either the liquid or solid fractions using the JEV G4 assay. However, effluent sample collected from the breeding shed on the same date revealed detection in the liquid fraction (3/3 positive with mean Cq: 34.6 ± 0.77) and 3/3 solid fraction replicates (mean Cq: 30.2 ± 0.08). Lab 2 also confirmed both samples as positive.

On 12/02/2025, effluent samples were collected from two locations (gestating shed and effluent pond) at Farm 6. At the outlet from the gestating shed, 1/3 liquid fraction sample tested positive (Cq: 37.2) for JEV using the Universal JEV assay, but no detection was recorded in Lab 2 using the JEV G4 assay. In the solid fraction samples from the gestating shed, 3/3 replicates were positive with a mean Cq value of 35.5 ± 0.40. Lab 2 confirmed the presence of JEV in these solid fraction samples, with 1/2 replicates positive (Cq 38.8). In the effluent from the pond, no JEV was detected in the liquid fraction samples, while the solid fraction samples showed a positive result in 2/3 replicates with a Cq value of 37.1 using the Universal JEV assay. However, Lab 2 did not detect JEV in these solid fraction samples using the JEV G4 assay. Veterinary cases in the herd were confirmed from samples that were collected from piglets on 31/01/2025, 12 days prior to the collection of effluent samples. Affected sows experienced abortion, stillbirth, fetal mummification, or gave birth to weak piglets. Reproductive failure can occur in sows infected with JEV up to approximately 60 to 70 days of gestation, while infection at later stages does not appear to affect piglets (28,29). Therefore, affected sows were likely infected approximately 6 to 8 weeks or more earlier, and were unlikely to be shedding at the time of effluent sampling, based on existing knowledge of virus shedding dynamics in pigs (18,19,21,22). In the absence of data confirming the persistence of infectious JEV we speculate that the positive effluent was residual virus shed during farrowing (e.g., from infected piglets and placental fluids), or from other infected sows experiencing an acute stage of infection.

Subsequent effluent samples collected from the gestating shed and effluent pond from Farm 6 on 21/02/2025, 24/02/2025, and 27/02/2025 were also JEV positive in Lab 1 using the JEV G4 assay. Lab 2 detected JEV in liquid fraction samples from the gestating shed and solid fraction of effluent pond samples on 21/02/2025, and in solid fraction of effluent pond on 24/02/2025 using the JEV G4 assay. Effluent sample (solid fraction) collected from Farm 6 gestating shed on 27/02/2025 were positive (1/3 replicates; Cq = 37.8), but liquid fraction was negative. Effluent sample (both liquid and solid fractions) from the pond was also positive by Lab 1. However, Lab 2 could not detect JEV in the effluent pond liquid fraction sample collected on 27/02/2025.

Interestingly, of the three environmental water samples collected from the borrow pit, one sample collected on 24/02/2025 was positive for JEV by both Lab 1 and 2 using the JEV G4 assay. This detection is significant because a borrow pit can collect stagnant water, creating an ideal breeding ground for mosquitoes, particularly *Culex* species, which are the primary vectors of JEV. Viruses present in or shed from infected larvae or present in infected mosquito eggs that were collected in these water samples may have been the source of the positive sample from the borrow pit. Subsequent environmental sample collected from the borrow pit was negative.

The RT-PCR results from Lab 1 and 2 revealed both agreements and discrepancies across different sample matrices and farm sampling locations (Table 1). For positive results in agreement, Cq values from Lab 1 testing were either comparable or lower than those from Lab 2. Ten samples also tested positive at Lab 1 but were negative (not detected) at Lab 2. The Cq values for these samples were all ≥36.6 and, for seven samples, only detected in 1/3 or 2/3 replicates at Lab 1. These discrepancies were attributed to a combination of differences in RT-PCR assay chemistry and sensitivity between the two laboratories, sub-sampling variability, samples with high Cq values at or near the ALOD, RNA degradation from freeze-thaw cycles, or sample degradation during shipment to Lab 2. To further validate the RT-PCR findings, the sequence obtained by the sanger sequencing was subjected to BLAST analysis, confirming 100% identity with JEV reference sequence OP904182. No mismatches or ambiguous bases were observed.

An important finding of this study is that solid fraction of effluent consistently yielded lower Cq values (i.e., higher concentrations of target nucleic acids) and a higher frequency of positive detections than liquid fraction effluent samples. This suggests that solid fraction of effluent may act as a more stable reservoir for JEV, supporting previous research indicating that viral particles preferentially adsorb to organic matter (27). Processing 50 mL of effluent samples was challenging due to high turbidity and the presence of suspended solids. To address this, we centrifuged the samples to separate the solid fraction from the liquid fraction before RNA extraction. The results suggest that analyzing only the solid fraction may be sufficient for JEV detection, potentially streamlining future surveillance efforts by reducing sample processing complexity.

## CONCLUSIONS

This study highlights the potential utility of effluent and environmental water surveillance for detecting JEV in piggery settings, with potential applications for emerging animal disease surveillance in a range of livestock industries. These findings provide a foundation for improving JEV monitoring strategies and supporting its mitigation in piggery operations. While this study focused on environmental sampling, specifically effluent from swine facilities, the findings suggest that such samples can serve as proxies for animal-level infection and may be integrated into broader One Health surveillance frameworks. Further research is needed to understand the dynamics and mode of virus shedding in infected pigs at different stages of production, particularly for infected pregnant sows for which there is a paucity of information reported. It is also important to investigate environmental factors such as temperature, pH, and effluent management practices that may influence the persistence of JEV in effluents. Additionally, assessing the correlation between viral detection in effluent compared to trapped mosquito pools, and oral fluids collected from piggeries could provide valuable insights into the most sensitive method or optimal combination of methods for cost-effective and efficient piggery-based surveillance for JEV. Continuous monitoring of farms is essential especially in summer (when environmental and ecological factors favour the transmission of mosquito-borne viruses) to determine whether virus is shed early before JEV outbreaks occur. In this study, we collected grab samples which were taken at a specific time. If JEV from infected pigs is intermittently excreted into effluents at a farm or shed level, a grab sample may not contain detectable levels of virus, leading to false negatives. Passive sampling may provide a more effective approach by continuous collection of samples over time, potentially increasing the sensitivity of viral detection and improving surveillance efficiency. Future studies should evaluate the feasibility and effectiveness of different passive sampling regimens for JEV monitoring using piggery effluent.

## ACKNOWLEDGEMENTS

We thank farm owners for providing samples.

## REFERENCES

1. Mackenzie JS, Williams DT. 2009. The zoonotic flaviviruses of southern, south-eastern and eastern Asia, and Australasia: the potential for emergent viruses. Zoonoses Public Health 56:338–356.

2. Mackenzie JS, Williams DT, van den Hurk AF, Smith DW, Currie BJ. 2022a. Japanese Encephalitis Virus: The emergence of genotype IV in Australia and its potential endemicity. Viruses 14:2480.

3. Liu Y, Smith W, Gebrewold M, Simpson SL, Williams DT, Wang X, Ahmed W. 2024. The effect of diurnal temperature fluctuations on the decay of Japanese Encephalitis and Murray Valley Encephalitis virus RNA seeded in piggery wastewater. ACS EST Water 4:4052–4060.

4. Mackenzie JS, Smith DW, Speers DJ. 2022b. Japanese encephalitis disease: overview of the virus, its risk to Australia and the need for better surveillance. Intern Med J 52:2029–2033.

5. van den Hurk AF, Ritchie SA, Mackenzie JS. 2009. Ecology and geographical expansion of Japanese encephalitis virus. Annu Rev Entomol. 54:17–35.

6. Hameed M, Wahaab A, Nawaz M, Khan S, Nazir J, Liu K, Wei J, Ma Z. 2021. Potential role of birds in Japanese Encephalitis Virus zoonotic transmission and genotype shift. Viruses 13:357.

7. Moore KT, Mangan MJ, Linnegar B, Athni TS, McCallum HI, Trewin BJ, Skinner E. 2025. Australian vertebrate hosts of Japanese encephalitis virus: a review of the evidence. Trans R Soc Trop Med Hyg 119:189–202.

8. Park SL, Huang Y-JS, Vanlandingham DL. 2022. Re-examining the importance of pigs in the transmission of Japanese Encephalitis Virus. Pathogens 11:575.

9. Khan A, Riaz R, Nadeem A, Amir A, Siddiqui T, Batool, UEA, Raufi N 2024. Japanese encephlu emergence in Australia: the potential population at risk. Ann. Med. Surg (Lond). 86:1540–1549.

10. Chen C, Wang Y, Kaur G, Adiga A, Espinoza B, Venkatramanan S, Warren A, Lewis B, Crow J, Singh R, Lorentz A, Toney D, Marathe M. 2024. Wastewater-based epidemiology for COVID-19 surveillance and beyond: A survey. Epidemics 49:100793.

11. Wolfe MK, Duong D, Shelden B, Chan EMG, Chan-Herur V, Hilton S, Paulos AH, Xu X-R S, Zulli A, White BJ, Boehm AB. 2024. Detection of hemagglutinin H5 Influenza A virus sequence in municipal wastewater solids at wastewater treatment plants with increases in Influenza A in Spring, 2004. Environ Sci Technol Lett. 11:526–532.

12. Hellmér M, Paxéus N, Magnius L, Enache L, Arnholm B, Johansson A, Bergström T, Norder H. 2014. Detection of pathogenic viruses in sewage provided early warnings of hepatitis A virus and norovirus outbreaks. Appl Environ Microbiol. 80:6771–6781.

13. Gupta P, Liao S, Ezekiel M, Novak N, Rossi A, LaCross N, Oakeson K, Rohrwasser A. 2023. Wastewater genomic surveillance captures early detection of Omicron in Utah. Microbiol Spectr 11:e0039123.

14. Xiao K, Zhang L. 2023. Wastewater pathogen surveillance based on One Health approach. Lancet Microbe 4:e297.

15. Ahmed W, Liu Y, Smith W, Ingall W, Belby M, Bivins A, Bertsch P, Williams DT, Richards K, Simpson S. 2024. Leveraging wastewater surveillance to detect viral diseases in livestock settings. Sci Total Environ 931:172593.

16. Fanok S, Monis PT, Keegan AR, King BJ. 2023. The detection of Japanese encephalitis virus in municipal wastewater during an acute disease outbreak. J Appl Microbiol 134:lxad275.

17. Lee WL, Gu X, Armas F, Leifels M, Wu F, Chandra F, Chua FJD, Syenina A, Chen H, Cheng D, Ooi EE, Wuertz S, Alm EJ, Thompson J. 2022. Monitoring human arboviral diseases through wastewater surveillance: Challenges, progress and future opportunities. Water Res 223:118904.

18. Ricklin ME, García-Nicolás O, Brechbühl D, Python S, Zumkehr B, Nougairede A, Charrel RN, Posthaus H, Oevermann A, Summerfield A. 2016. Vector-free transmission and persistence of Japanese encephalitis virus in pigs. Nat Commun 7:10832.

19. Lyons AC, Huang YS, Park SL, Ayers VB, Hettenbach SM, Higgs S, McVey DS, Noronha L, Hsu WW, Vanlandingham DL. 2018. Shedding of Japanese Encephalitis virus in oral fluid of infected swine. Vector Borne Zoonotic Dis. 18:469–474.

20. Chiou SS, Chen J-M, Chen Y-Y, Chia M-Y, Fan Y-C. 2021. The feasibility of field collected pig oronasal secretions as specimens for the virologic surveillance of Japanese encephalitis virus. PLoS Negl Trop Dis 15:e0009977.

21. Hick PM, Finlaison DS, Parrish K, Gu X, Hayton P, O’Connor T, Read A, Zhang J, Spiers ZB, Pinczowski P, Ngo AL, Kirkland PD. 2024. Experimental Infections of Pigs with Japanese Encephalitis Virus Genotype 4. Microorganisms 12:2163.

22. Cool K. 2020. Characterizing the fecal shedding of swine infected with Japanese encephalitis virus. Master Dissertation, Kansas State University, Manhattan, Kansas. Available from: https://krex.k-state.edu/bitstream/handle/2097/40926/KonnerCool2020.pdf. Retrieved 11 Mar 2024

23. ProMED. 2025. Japanese encephalitis - Australia (02): (VI, NS) spread. Archive number 20250114.8721340.

24. Akter J, Smith WJM, Liu Y, Kim I, Simpson SL, Thai P, Korajkic A, Ahmed W. 2024. Comparison of adsorption-extraction (AE) workflows for improved measurements of viral and bacterial nucleic acid in untreated wastewater. Sci Total Environ 908: 167966.

25. Shao N. Li F, Nie K, Fu SH, Zhang WJ, He Y, Lei WW, Wang QY, Liang GD, Cao YX, Wang HU. 2018. TaqMan real-time RT-PCR assay for detecting and differentiating Japanese Encephalitis Virus. Biomed Environ Sci 31:208–214.

26. Staley C, Gordon KV, Schoen ME, Harwood VJ. 2012. Performance of two quantitative PCR methods for microbial source tracking of human sewage and implications for microbial risk assessment in recreational waters. Appl Environ Microbiol 78:7317–7326.

27. Roldan-Hernandez L, Van Oost C, Boehm AB. 2025. Solid–liquid partitioning of dengue, West Nile, Zika, hepatitis A, influenza A, and SARS-CoV-2 viruses in wastewater from across the USA. Environ. Sci.: Water Res. Technol. 11:88–99.

28. Williams DT, Mackenzie JS, Bingham J. 2019. Flaviviruses. In Diseases of Swine; Wiley: Hoboken, NJ, USA, 530–543.

29. Platt KB, Joo HS. 2006. Japanese encephalitis and West Nile viruses. In: Diseases of Swine, 9th edition. Straw BE, Zimmerman JJ, D’Allaire S, Taylor DJ (editors). Blackwell Publishing, Ames, Iowa, pp. 359–365.

30. Chapagain S, Pal Singh P, Le K, Safronetz D, Wood H, Karniychuk, U. 2022. Japanese encephalitis virus persists in the human reproductive epithelium and porcine reproductive tissues. PLoS Negl Trop Dis 16:e0010656.

